# Different mechanisms drive the maintenance of polymorphism at loci subject to strong versus weak fluctuating selection

**DOI:** 10.1101/164723

**Authors:** Jason Bertram, Joanna Masel

## Abstract

The long-running debate about the role of selection in maintaining genetic variation has been given new impetus by the discovery of hundreds of seasonally oscillating polymorphisms in wild *Drosophila,* possibly stabilized by an alternating summer-winter selection regime. Historically there has been skepticism about the potential of temporal variation to balance polymorphism, because selection must be strong to have a meaningful stabilizing effect — unless dominance also varies over time (“reversal of dominance”). Here we develop a simplified model of seasonally variable selection that simultaneously incorporates four different stabilizing mechanisms, including two genetic mechanisms (“cumulative overdominance” and reversal of dominance), as well as ecological “storage” (“protection from selection” and boom-bust demography). We use our model to compare the stabilizing effects of these mechanisms. Although reversal of dominance has by far the greatest stabilizing effect, we argue that the three other mechanisms could also stabilize polymorphism under plausible conditions, particularly when all three are present. With many loci subject to diminishing returns epistasis, reversal of dominance stabilizes many alleles of small effect. This makes the combination of the other three mechanisms, which are incapable of stabilizing small effect alleles, a better candidate for stabilizing the detectable frequency oscillations of large effect alleles.

## 1 Introduction

Understanding how populations maintain genetic variation for fitness-associated traits is a foundational problem in evolutionary biology. In the “classical view”, genetic variation appears by mutation and is removed by a combination of selection and random genetic drift, resulting in unstable variation that continually turns over. In contrast, the “balance view” posits that genetic variation is stably maintained by selection.

The theoretical basis of mutational variation is simple: mutations will maintain variation if mutation rates are sufficiently high. Empirically, deleterious mutation rates can sometimes exceed one per individual per generation (Lynch et al., 1999; Lesecque et al., 2012), and it is well established that transient mutations contribute substantially to standing genetic variation in fitness-associated traits (Keightley and Halligan, 2008; Charlesworth, 2015).

The contribution of balancing selection to standing genetic variation can also be substantial (Bitarello et al., 2018; Good et al., 2017; Charlesworth, 2015). In a particularly striking example, sequencing studies in temperate *Drosophila* populations have found hundreds of polymorphic loci which undergo seasonal oscillations in allele frequency (Bergland et al., 2014; Machado et al., 2018). Many of these polymorphisms appear to be ancient (Bergland et al., 2014), suggesting pervasive, long-term balancing selection.

The possibility of pervasive balancing selection poses a theoretical problem. Unlike the case of mutational variation, the conditions leading to balanced polymorphism are not resolved (Yi and Dean, 2013; Svardal et al., 2015; Wittmann et al., 2017). Fundamentally, stable polymorphism requires negatively frequency-dependent selection, i.e. low frequency alleles must be favored over high frequency alleles. In temporally and spatially uniform environments, instances of negative frequency-dependence are largely restricted to immune system genes (Aguilar et al., 2004), Batesian mimicry (Kunte, 2009) and self incompatibility loci (Charlesworth, 2006). Heterogeneous selection substantially broadens the types of loci potentially subject to negative frequency-dependence, since the latter only needs to arise when averaged over the environmental heterogeneity, rather than within each generation, or within spatially homogeneous subpopulations (Frank and Slatkin, 1990; Svardal et al., 2015). The theoretical problem is to explain how environmental heterogeneity can lead to sufficiently strong balancing selection under a plausible range of conditions.

In the *Drosophila* example above, polymorphism appears to be stabilized by temporal heterogeneity, specifically adaptation to alternating seasonal selection regimes (Bergland et al., 2014; Machado et al., 2018). Inspired by this example, we restrict our attention to temporal heterogeneity (for discussion of spatial heterogeneity see Frank and Slatkin (1990); Hedrick (2006); Svardal et al. (2015) and references therein). We distinguish between two broad classes of mechanism by which temporal heterogeneity can stabilize polymorphism.

The first are mechanisms which rely on purely genetic factors such as diploid Mendelian inheritance and epistasis. Early studies of fluctuating selection focused on the consequences of incomplete dominance. Rare alleles all appear in heterozygotes, and so incomplete dominance can reduce selection against rare alleles in the environments they are unsuited to, making selection negatively frequency-dependent (Dempster, 1955; Haldane and Jayakar, 1963). Following Dempster (1955), we call this “cumulative overdominance”.

“Reversal of dominance” (also known as “segregation lift”; Wittmann et al. 2017) also relies on diploid Mendelian inheritance, but additionally requires dominance to change over time such that the favored allele at any moment is dominant. Reversal of dominance was the basis of Gillespie’s SAS-CFF model of non-neutral genetic variation (Gillespie, 1978; Hedrick, 1986). More recently, Wittmann et al. (2017) showed that reversal of dominance can simultaneously stabilize hundreds of polymorphic loci in mutation-selection balance, without an attendant genetic load problem (i.e. the large number of segregating loci did not imply fitness differences so large that they require an implausibly large reproductive capacity; see also Gillespie 2010, pp. 74).

Sign epistasis between locus pairs can stabilize polymorphism via an interaction with linkage disequilibrium (Novak and Barton, 2017). While this mechanism may be important at some loci, we will not discuss it further because it is not clear how it should be applied to more than two loci (other than that there are simply many pairs of sign-epistatic loci).

Our second class of stabilizing mechanisms are ecological i.e. they depend on ecological interactions between individuals rather than genetic interactions between alleles. We consider two mechanisms in particular, which are both particular instances of the “storage effect”. The storage effect encompasses a broad class of fluctuation-dependent coexistence scenarios with potentially very different biological underpinnings (Chesson, 2000). Historically, the storage effect was introduced using a model with two life stages, adults and juveniles, with overlapping generations and competition among juveniles only (Chesson and Warner, 1981). In this model, some adults are “stored” over multiple rounds of juvenile recruitment without themselves experiencing competition. This reduces losses in unfavorable environments, and allows rare genotypes to rapidly recoup their losses when favorable environmental conditions return. Overlapping generations and similar phenomena like resting eggs in *Daphnia* have so far been the focus of genetic applications of the storage effect (Ellner and Hairston 1994; Yi and Dean 2013; Hedrick 2006; Svardal et al. 2015; although see Gulisija et al. 2016 for an epistatic interpretation of the storage effect). What ties these phenomena together is “protection from selection”, that is, a fraction of each genotype/species experiences weakened selection. This is the first of our ecological mechanisms.

The storage effect also includes scenarios where there is no protection from selection in the above sense. We will consider a stabilization phenomenon that occurs when population density undergoes repeated “boom-bust” demographic cycles (Yi and Dean, 2013; Li and Chesson, 2016), as is the case in temperate *Drosophila* populations driven in part by seasonal variation in fruit availability (Bergland et al., 2014). This phenomenon is a consequence of crowding effects at higher densities, which can result in longer generation times. When a rapid summer grower genotype is common, the population will reach high densities more rapidly than when a slow-growing winter-adapted type is common, resulting in fewer versus more generations of selection in the summer for the two respective cases. This creates frequency-dependent selection for the seasonal cycle as a whole, which can stabilize polymorphism (Yi and Dean, 2013).

The preceding paragraphs demonstrate that there is no shortage of mechanisms that can stabilize polymorphism in the presence of temporal variability. Nevertheless, temporal variability has not been regarded as a robust basis for arguing that balanced polymorphism should be prevalent, or for explaining known balanced polymorphisms, even though temporal variability is ubiquitous. The reason for this is that most stability mechanisms based on temporal variability require fluctuating selection to be strong in order to have an appreciable stabilizing effect (Hoekstra et al., 1985; Smith and Hoekstra, 1980; Hedrick, 1986, 2006). That is, selection must strongly favor different genotypes at different times to be able to counteract a non-negligible difference in time-averaged genotypic fitness. For example, taken in isolation, cumulative overdominance is too weak to stabilize polymorphism unless selection coefficients are at least of order 0.1 (Hoekstra et al., 1985, Fig. 3). By comparison, in the above *Drosophila* example the overall absolute change in allele frequency over a season — which constitutes multiple generations — is itself only of order ~ 0.1, implying smaller selection coefficients (Wittmann et al., 2017; Machado et al., 2018).

Reversal of dominance is the major exception to the strong-selection requirement, and has recently been investigated as a possible candidate to explain multi-locus balancing selection in *Drosophila* (Wittmann et al., 2017). However, escaping the strong selection requirement means that reversal of dominance can stabilize alleles of weak effect, which poses different problems. Under the empirically well-supported assumption of diminishing-returns epistasis (Chou et al., 2011; Kryazhimskiy et al., 2014), Wittmann et al. (2017) found that the majority of polymorphic loci in mutation-selection-drift balance had fitness effects so small that their frequency oscillations were undetectable — at most ~ 10 detectably oscillating loci could be stabilized. Because reversal of dominance is able to stabilize polymorphism between segregating alleles with weak fitness effects, the model predicted the accumulation of weak-effect polymorphic loci. In the presence of diminishing returns epistasis, this accumulation made the model incapable of generating stable polymorphisms of sufficiently strong effects to match the many detectably oscillating loci that have been observed in *Drosophila*. We are therefore faced with the conundrum that most of the stability mechanisms dependent on temporal variability require selection that is implausibly strong, but the more effective reversal of dominance mechanism allows selection to be so weak as to create other difficulties (it is also unknown whether reversal of dominance is likely to be prevalent in nature; see Gillespie 1978 and Wittmann et al. 2017 for discussion).

Here we argue that ecological coexistence mechanisms help to explain how it is possible to have a large number of detectably oscillating polymorphisms stabilized by temporal variability. The ecological mechanisms we consider also have a strong selection requirement, offering nothing new over and above cumulative overdominance in this respect. However, we will show that the ecological mechanisms operate in conjunction with cumulative overdominance. This expands the region of allelic fitness effects compatible with balanced polymorphism, partly mitigating the strong selection requirement. Moreover, this mitigation occurs in such a way that the diminishing returns problem that arises with reversal of dominance (discussed above) is avoided. Together these results give a more favorable view of the stabilizing potential of temporal variability, and could help to explain the *Drosophila* findings of Bergland et al. (2014) and Machado et al. (2018).

Our analysis rests on a simplified model of fluctuating selection that incorporates reversal of dominance, cumulative overdominance, protection from selection, and boom-bust demography all at once. We assume that the environment alternates between two states, “summer” and “winter”. This significantly simplifies the mathematical analysis compared to the more general case of a stochastic environment (e.g. Chesson and Warner 1981). We use our model to derive a combined stability condition including the effects of all of these mechanisms, which reveals both their individual contributions to stability, as well as their interactions.

## 2 Basic model and assumptions

Our basic model is a straightforward extension of the standard equations for selection in diploids. We assume that a winter allele W and a summer allele S are segregating at one locus in a randomly mating population. Each iteration of the model involves one round of random mating and juvenile recruitment to reproductive maturity. Some fraction of the copies of each allele are completely “protected” from selection, so that the allele frequencies within the protected fraction remain constant from one iteration to the next. The classic example of this form of protection is generational overlap (Chesson and Warner, 1981): if only a fraction 1 – *f* of reproductively mature individuals die and are replaced each round of juvenile recruitment, then alleles in the remaining fraction *f* experience no viability selection. Note that the particular copies of each allele that are protected may change between iterations. The environment alternates between summer and winter such that the per-iteration relative fitnesses of unprotected alleles take the values in Table 1. The change in winter allele frequency p_W_ over one iteration is thus given by

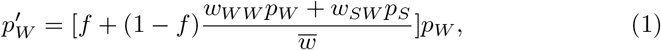

and similarly for the summer allele, where 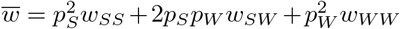 is the mean relative fitness of unprotected alleles. If there is no protection (*f* = 0), we have the usual 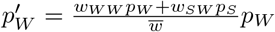.

**Table 1:** Relative fitness values for unprotected summer/winter alleles in an alternating summer/winter environment. The variables *a_S_* and *a_W_* denote the juvenile advantages of the *S* and *W* alleles in the summer and winter respectively.

|  | Summer homozygote | Heterozygote | Winter homozygote |
| --- | --- | --- | --- |
| Summer | $w_{SS}(S) = 1 + a_S$ | $w_{SW}(S) = 1 + d_S a_S$ | $w_{WW}(S) = 1$ |
| Winter | $w_{SS}(W) = 1$ | $w_{SW}(W) = 1 + d_W a_W$ | $w_{WW}(W) = 1 + a_W$ |

For polymorphism between *W* and *S* to be stable, both *W* and *S* alleles must increase in frequency over a full summer/winter cycle when rare. We will generally assume that the winter allele is rare (*p_W_* ≪ 1); the behavior of a rare summer allele will then be obtained by swapping the *S* and *W* labels. Since *W* is rare, 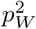 terms are negligibly small in Eq. (1), which thus simplifies to

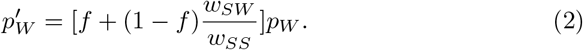

Eq. (2) expresses the fact that almost all individuals are *S* homozygotes, and almost all copies of *W* occur in heterozygotes.

We denote number of summer and winter iterations by *i_S_* and *i_W_* respectively. Therefore, the condition for the rare winter allele to increase in frequency over a complete seasonal cycle is

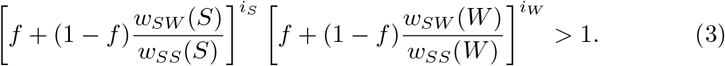

Eq. (3) will be the foundation for much of the analysis presented below. When we come to consider boom-bust population demography we will also evaluate some extensions of Eq. (3) that are better equipped for handling the effects of non-constant population density.

## 3 Results

### 3.1 Stabilizing mechanisms

In this section we compare four mechanisms that can stabilize polymorphism at a single locus in the presence of fluctuating selection: cumulative overdominance, reversal of dominance, protection from selection, and boom-bust population demography. We start by considering the first three of these separately, since these are substantially simpler to analyze in the absence of boom-bust demography. In particular, we can make the simplifying assumption that *i_S_* and *i_W_* are constants (that is, in each season selection proceeds in the same way for a fixed number of iterations; as we will see, this is not the case in the presence of boom-bust demography).

To further simplify, we begin by setting *i_S_* = *i_W_* (unequal but constant values of *i_S_* and *i_W_* do not induce frequency-dependence and thus do not stabilize polymorphism). It is then straightforward to show that Eq. (3) becomes (details in Appendix A)

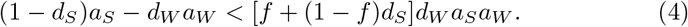

Below we use special cases of Eq. (4) to isolate the effects of cumulative overdominance, reversal of dominance, and protection from selection. We then extend our analysis to incorporate the effects of boom-bust population demography.

#### 3.1.1 Cumulative overdominance and reversal of dominance

We first consider the case of no protection (*f* = 0) in Eq. (4). Polymorphism can then be stabilized in two distinct ways. First, cumulative overdominance (Dempster, 1955) is a consequence of weakened negative selection on rare winter alleles in the summer due to incomplete dominance. Its effect can be seen most easily by assuming that dominance is constant between seasons, such that the winter dominance is the complement of the summer dominance *d_W_* = 1 – *d*_S_.

Eq. (4) then simplifies to

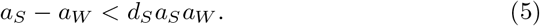

Eq. (5) shows that the rare winter allele can persist even if the summer allele is superior (*a_S_* > *a_W_*). The stabilizing effect gets stronger with increasing summer dominance *d_S_*. Strong fluctuating selection is needed for stability, since the product of juvenile advantages *a_W_a_S_* appears on the right hand side (the asymmetry between allelic fitness effects can be at most second order in the juvenile advantages *a_S_* and *a_W_*).

Combining Eq. (5) with the corresponding condition for a rare summer allele (given by Eq. (5) with the S and W labels swapped) defines a region of stable polymorphism for a given dominance (Fig. 1a, red). The requirement for strong fluctuating selection can be seen not just by the small size of the red region as a whole, but more specifically by the way it tapers to little more than a line for small values of *a_S_* and *a_W_*.

**Figure 1:**
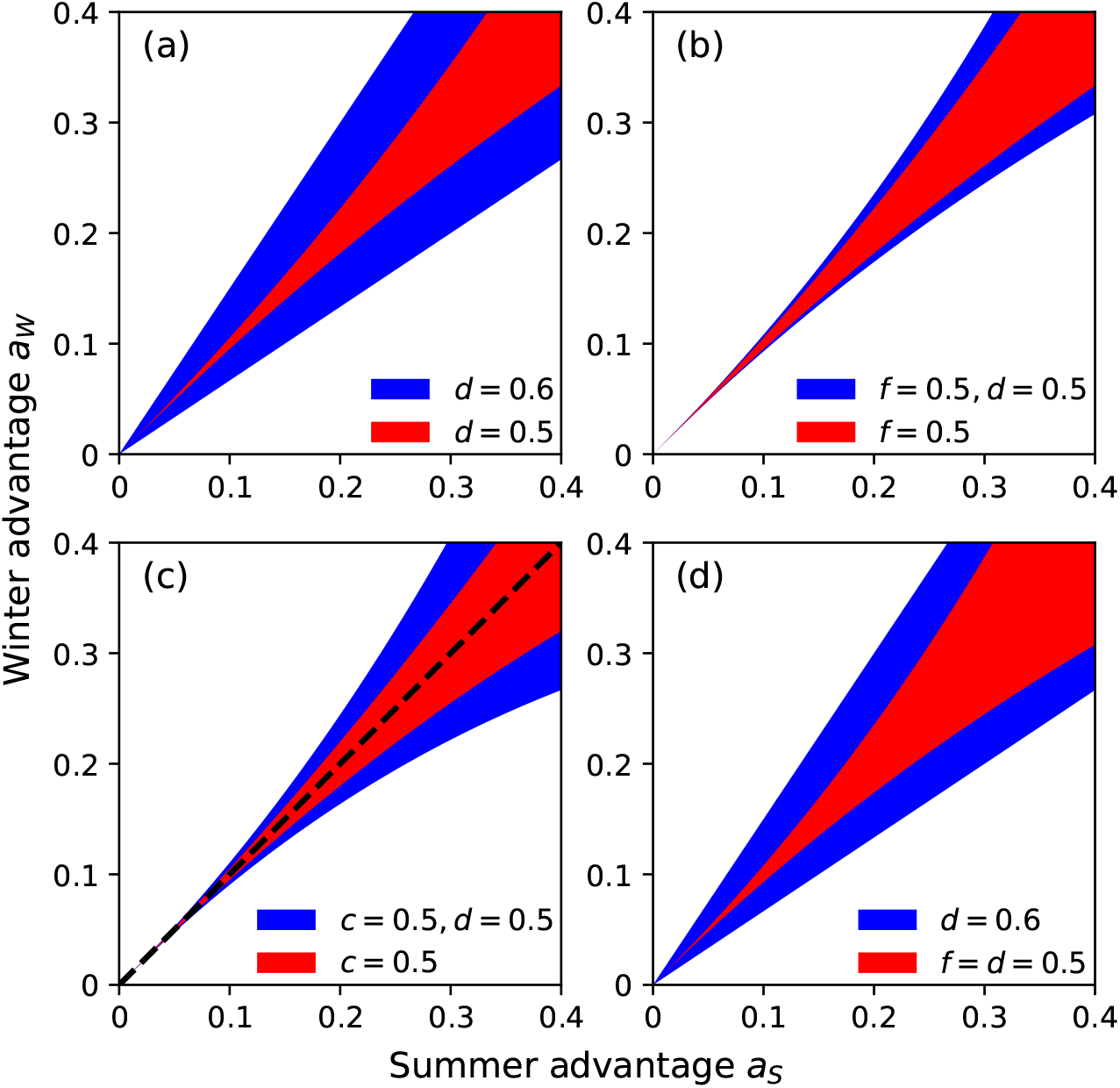
(a) Cumulative overdominance alone only stabilizes a small region (red; *d_S_* = *d_W_* = 0.5 in Eq. (5)), especially if selection does not fluctuate strongly (*a_S_* and *a_W_* are not large). Reversal of dominance creates a much larger region of stability (blue; *d* = 0.6 in Eq. (7)). (b) Protection from selection in haploids (Eq. (8)) stabilizes a region with the same shape as cumulative overdominance in diploids, and magnitude set by *f*. In diploids with both cumulative overdominance and protection from selection, the stabilized region is only modestly larger than protection from selection in haploids (blue; *d_S_* = *d_W_* = 0.5 and *f* = 0.5 in Eq. (4)). (c) Boom-bust demography alone (red; *c* = 0.5 in Eq. (11)) is similar to cumulative overdominance, but is not symmetric (more of the red lies below than above the dashed line) since it hinges on the summer advantage *a_S_*. (d) Reversal of dominance alone (blue; Eq. (7)) is effective at stabilizing alleles of any strength, but the combination of cumulative overdominance and protection from selection (red) is similarly effective for larger effect alleles.

The second way that stable polymorphism can occur is reversal of dominance. This requires dominance to alternate between seasons. For simplicity, let the magnitude of alternating dominance be constant, i.e. *d_S_* = *d_W_* = *d*. Then Eq. (4) can be written as

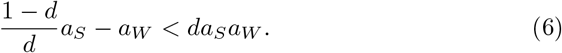

The right hand side of Eq. (6) represents the effects of cumulative overdominance (compare Eq. (5)). The effects of reversal of dominance can be viewed in isolation by setting this term to zero,

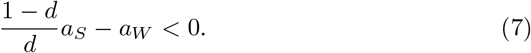

In Eq. (7), the summer advantage *a_S_* is multiplied by the factor (1 – *d*)/*d*, which is smaller than 1 if the favored allele is dominant 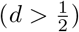. This stabilizes polymorphism by partly offsetting any advantage that the common summer allele may have. Unlike cumulative overdominance, which is second order in the juvenile advantages, reversal of dominance has a first order effect (Fig. 1a, blue). This powerful effect arises because rare alleles are more often found in heterozygotes, thus granting them the major benefit that heterozygotes are always closer in fitness to the favored homozygote than to the unfavored homozygote.

#### 3.1.2 Protection from selection

We now examine the effects of protection from selection, allowing *f* > 0 (but still keeping the number of summer and winter iterations equal *i_S_* = *i_W_*). We do this in two stages for ease of comparison with the diploid case discussed above. First, we consider a haploid variant of our model where the winter genotype has *w_W_*(*S*) = 1 and *w_W_*(*W*) = 1 + *a_W_*, and the summer genotype *w_S_*(*S*) = 1 + *a_S_* and *w_S_*(*W*) = 1. It is then easily verified that the condition for persistence of a rare winter allele is

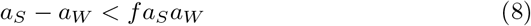

This condition can be derived from scratch in a similar way to Eq. (4). Alternatively, we can simply observe that the haploid case is a special instance of the diploid model where *d_S_* = 0 and *d_W_* = 1 when the winter allele is rare, and *d_S_* = 1 and *d_W_* = 0 when the summer allele is rare (compare the haploid fitnesses with Table 1, and then substitute these values into Eq. (4)).

Eq. (8) is the same condition as Eq. (5), but with *d_S_* replaced by *f*. Thus, as in the case of cumulative overdominance, stable polymorphism under protection from selection requires fluctuating selection to be strong (Fig. 1b, red). Intuitively, protection favors rarity because, in the absence of protection, an abundant type can easily displace a large fraction of a rare type’s individuals when the abundant type is favored, due to sheer numerical advantage. Protection thus limits the rare type’s fractional losses. By contrast, a rare type can only displace a tiny fraction of an abundant type’s individuals when the rare type is favored, and its growth is therefore not limited by protection of the abundant type.

In diploids, the effect of protection from selection is represented by the *f* + (1 – *f*) *d_S_* factor on the right hand side of Eq. (4), representing an increase of (1 – *f*)*d_S_* over the corresponding factor in the haploid case Eq. (8). This increase gets smaller for larger f, because cumulative overdominance only affects unprotected allele copies.

#### 3.1.3 Boom-bust demography

We now derive a condition analogous to Eq. (4) that accounts for an additional source of negative frequency dependence induced by boom-bust demography. This boom-bust demography stabilizing mechanism requires: (1) alternating periods of population expansion and collapse; (2) selection favoring the S allele in the summer must be weaker at higher population density; (3) the overall amount of growth in the summer (as measured by the growth factor *D* from summer start to summer end) must be frequency-independent. We comment on the plausibility of (2) and (3) at the end of this section; first we illustrate the essential features of the boom-bust demography stabilizing mechanism by considering an idealized scenario in which selection ceases entirely at high density (our approach follows Yi and Dean (2013)).

Although our basic model Eq. (1) does not explicitly represent population density, it can nevertheless be used to illustrate the boom-bust stabilizing mechanism by relaxing our previous assumption that *i_S_* is constant. To show why this is so, we first assume for simplicity that there is no protection (the interaction between protection and boom-bust demography is addressed in the following section). Next, we assume that each genotype grows at a constant rate in the summer (*r*-selection). These genotype-specific absolute growth rates are assumed to be proportional to the relative fitnesses in Table 1, and will be denoted by r. For instance, the abundance of a rare winter allele grows according to 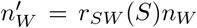, while 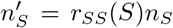 for the summer allele, where *r_SW_*(*S*) = (1 + *d_S_a_S_*)*r_WW_*(*S*) and *r_SS_*(*S*) = (1 + *a_S_*)*r_WW_*(*S*). Density-dependence is incorporated by assuming that total population density N reaches carrying capacity before the end of summer, at which point selection ceases entirely (satisfying condition (2)).

The winter is assumed to consist of density-independent mortality at a constant rate α such that, in the rare-*W* case, 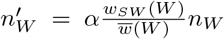 and 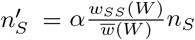 where *α* < 1 and *w* is defined in Table 1. Ecologically, this could represent the gradual loss of viable overwintering sites, with the winter allele being better at securing and holding those sites. Since the population growth factor *D* in the summer is equal to the winter decline when the boom-bust cycle is in a steady state, this winter mortality model ensures that *D* is frequency-independent such that it has the same value in the rare-*S* and rare-*W* cases.

The allele frequencies in the above model obey Eq. (1) with *f* = 0, and so Eq. (3) applies with *f* = 0. However, the number of iterations of selection in the summer *i_S_* is not constant in Eq. (3), as was assumed in the preceding sections. Now *i_S_* depends on how many iterations of growth it takes for the population to reach carrying capacity. This induces negative frequency-dependence because the common type determines the mean population growth rate. Thus, a lower winter allele frequency implies more rapid population saturation, and hence fewer iterations of negative selection experienced by the winter allele. Conversely, a lower summer allele frequency implies more iterations of positive selection on the summer allele (Fig. 2). The result is a stabilizing effect.

**Figure 2:**
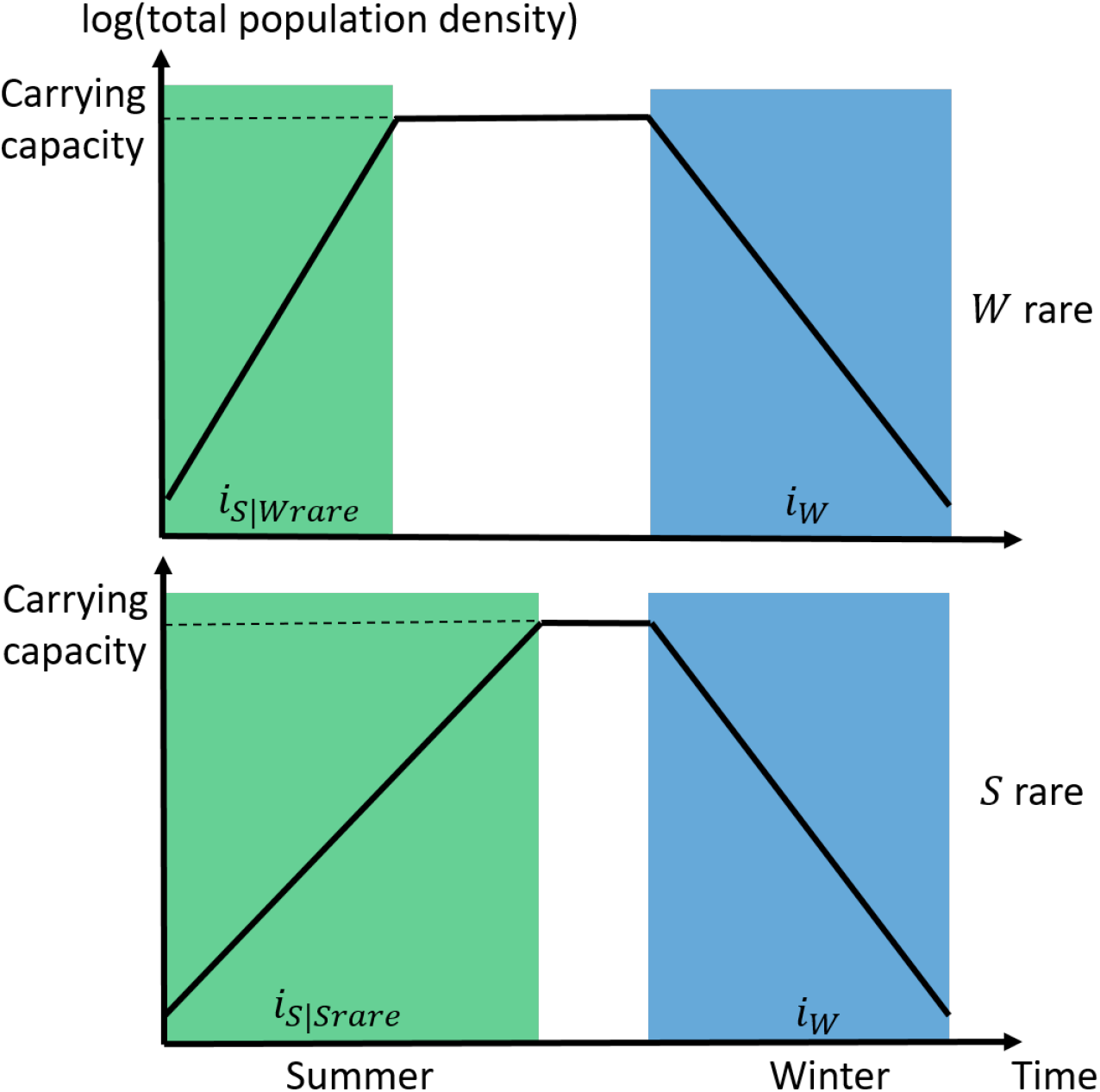
Illustration of a winter/summer boom-bust demographic cycle in a simplified model of exponential growth up to saturation in the summer, and exponential decline in the winter. The periods of selection in the summer and winter are marked with green and blue boxes respectively. Selection ceases when total population density is at carrying capacity, causing there to be more iterations of summer selection when the summer allele is rare.

In Appendix B we use Eq. (3) to quantify the boom-bust stabilizing effect under the above assumptions. The number of summer iterations *i_S_* is shown to depend on frequency as follows

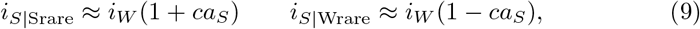

where *c* is a constant approximately equal to 1/2. The difference in *i_S_* between the summer-rare versus winter-rare frequency regimes is therefore first order in the summer advantage *a_S_*. This introduces second order terms into the stability condition Eq. (4), giving

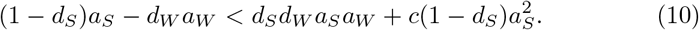

The condition for a rare summer allele to grow is obtained as before by swapping labels in all terms apart from the last, which is replaced by 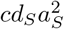 (details in Appendix B). Similar to our analysis of protection from selection, the effect of boom-bust demography can be isolated by considering the haploid case (*d_S_* = 0 and *d_W_* = 1 when the winter allele is rare, *d_S_* = 1 and *d_W_* = 0 when the summer allele is rare), which gives

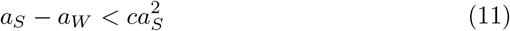

for a rare winter allele (and 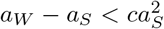 for rare *S*).

Eq. (10) and (11) show that the strength of the boom-bust stabilizing mechanism is second order in the juvenile advantages, like cumulative overdominance and protection from selection, but with the difference that the stabilizing term depends on the summer advantage only (Fig. 1c, red). Eq. (10) also shows that there is no interaction between the stabilizing contributions of boom-bust demography and cumulative overdominance (the corresponding terms on the right hand side are simply added). Intuitively we do not expect dominance to affect the boom-bust mechanism because the latter depends on the growth rate of *W* and *S* homozygotes; these determine the overall population growth rate when the corresponding allele is common.

The above model of constant growth followed by stasis at carrying capacity follows Yi and Dean’s (2013) model of coexistence in a microbial serial transfer experiment. The alternating “seasons” in Yi and Dean (2013) are both booms, each followed by a selectively neutral dilution step by a fixed dilution factor *D*. Here the environmental cycle only involves a single boom followed by a non-neutral bust, and we have shown that a rarity advantage can also arise in this case under conditions (1)-(3).

Are conditions (2) and (3) plausible? The density-dependent selection in (2) could reflect trade-offs between traits favorable at low vs high densities, but (2) could also occur simply because less selection occurs at high density compared to the initial low-density “boom” (e.g. due to reduced population turnover). Condition (3) is open to the objection that we might expect total population size at the end of the winter to be smaller if the winter survivor *W* is rare than if it is common (Fig. 2). In this case the summer growth factor D up to carrying capacity would itself depend on frequency in a way that eliminates the rarity advantage induced by density-dependent growth (there would be no negative frequency dependence if we had assumed *r*-selection in the winter as well). Thus, the plausibility of condition (3) hinges on the extent to which selection in the winter represents simple differences in extrinsic mortality rates versus a relatively inflexible population bottleneck (e.g. scarce overwintering sites).

To complete our analysis of the boom-bust stabilizing mechanism, we supplement the above proof of principle with a simulation example that incorporates a more realistic description of density-dependence. To preclude the possible stabilizing effects of diploidy discussed in the preceding sections, we restrict ourselves to the haploid case. We assume that growth in the summer follows the Ricker model 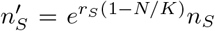 and 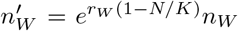 where *r_S_* = (1 + *a_S_*)*r_W_* and *N* = *n_S_* + *n_W_*. This model ensures that selection favoring the *S* allele gets weaker at higher densities (condition (2) above). Our motivation for using the Ricker model is that the comparably simple discrete-time logistic model 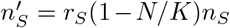 does not satisfy condition (2) because selection is density-independent (unless we additionally assume that *K* differs between alleles; this is both more complicated and potentially introduces coexistence in a constant environment which would make analysing the boom-bust stabilizing mechanism impossible). Winter mortality is the haploid version of the model used above to obtain Eq. (10). Fig. 3 shows the frequency and abundance trajectories of *S* and *W* haplotypes coexisting stably due to the boom-bust demography coexistence mechanism.

**Figure 3:**
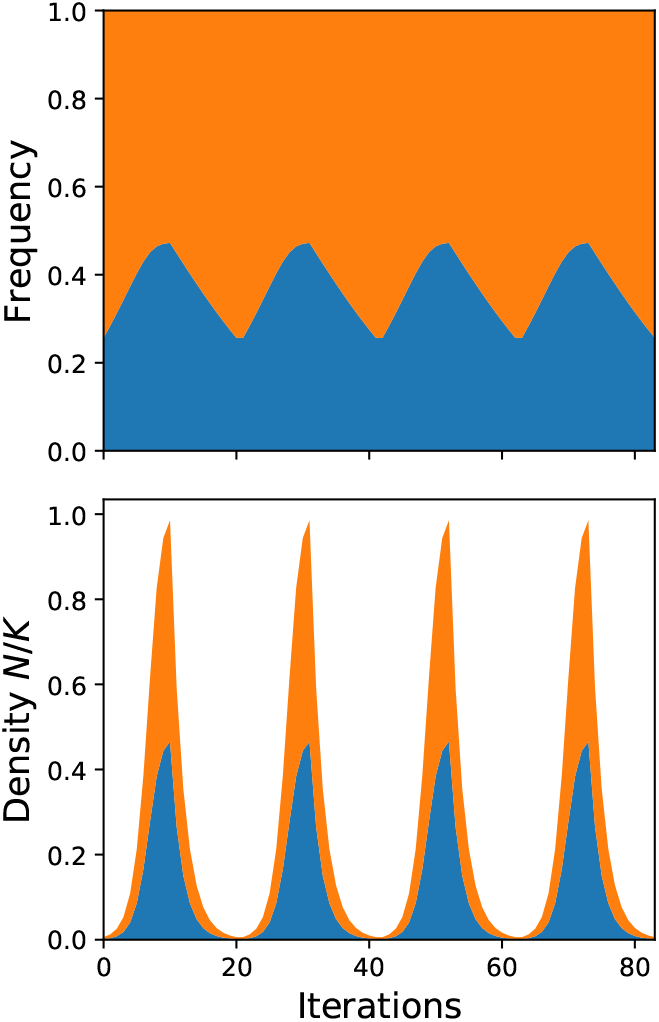
An example of haploid polymorphism stabilized by boom-bust demography where selection in the summer follows the Ricker model 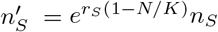 and winter mortality occurs at a constant rate 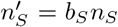 (blue areas indicate the summer allele). Parameters: *α* = 0.6, *a_W_* = 0.1, *r_S_* = 0.7, *a_S_* = 0.2, *K* = 10^5^, 10 generations/season.

#### 3.1.4 Combining protection and boom-bust demography

In the previous section we gave a simplified treatment of the boom-bust stabilizing mechanism, which assumed (among other things) that there was no protection. Here we consider the combined effects of protection and boom-bust demography.

In addition to allowing the boom-bust stabilizing mechanism to occur, boom-bust demographic cycles can also modify the stabilizing effects of protection, even if the boom-bust stabilizing mechanism is not present (e.g. because selection favoring the summer allele in the summer does not get weaker with increasing density). The reason is that the effect of protecting a fraction f of the allele copies in a given iteration depends on how many juveniles are recruited in the next iteration. There will be essentially no protective effect if the protected alleles are swamped by juveniles during a demographic boom, and conversely the protective effect is amplified in a shrinking population. To illustrate this point, consider a simple extension of Eq. (2) which allows population density to vary depending on the magnitude of juvenile recruitment:

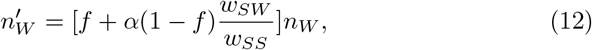

and similarly for a rare summer allele. Here *α* represents the magnitude of juvenile recruitment, and takes the constant values *α_S_* > 1 in the summer and *α_W_* < 1 in the winter (our original model Eq. (2) is retrieved when *α_S_* = *α_W_* = 1, corresponding to constant total population size *N*). If the population is expanding rapidly (*α_S_* ≫ 1), then the f term in Eq. (12) will be negligibly small by comparison and it will be as though there is no protection in the summer. Conversely, the influence of the f term will be amplified in the winter (when *α_W_* < 1) compared to Eq. (2).

To measure the stabilizing effect in Eq. (12), we calculate the average of each allele’s growth when rare over a full seasonal cycle,

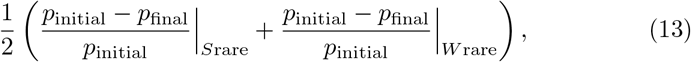

where the juvenile advantages *a_S_* = *a_W_* are assumed to be equal. Eq. (13) allows us to quantify stabilizing effects numerically when we do not have neat analytical results such as Eq. (4). Importantly, since we average over the two rarity cases, Eq. (13) is not sensitive to any asymmetries between winter and summer that might intrinsically favor the S or W allele even though *a_S_* = *a*_W_. Such asymmetries could be confused for a rarity advantage/disadvantage if only one of the rarity scenarios is examined in isolation, and are difficult to avoid in all but the simplest models (e.g. Eq. (2)). The stabilizing effect of protection in Eq. (12), as measured by Eq. (13), can be enhanced or diminished relative to the constant-N case depending on the values of α_S_, α_W_ and f. The effect is enhanced in many cases of biological relevance; Fig. 4 shows two examples (solid and black dashed lines). Intuitively, the reason for this is that the enhanced protective effect due to the winter bust *α_W_* < 1 expands the overall region of coexistence more than the reduced protective effect due to the summer boom truncates the region of coexistence – but the region of coexistence will be asymmetric (see Appendix C for more details).

**Figure 4:**
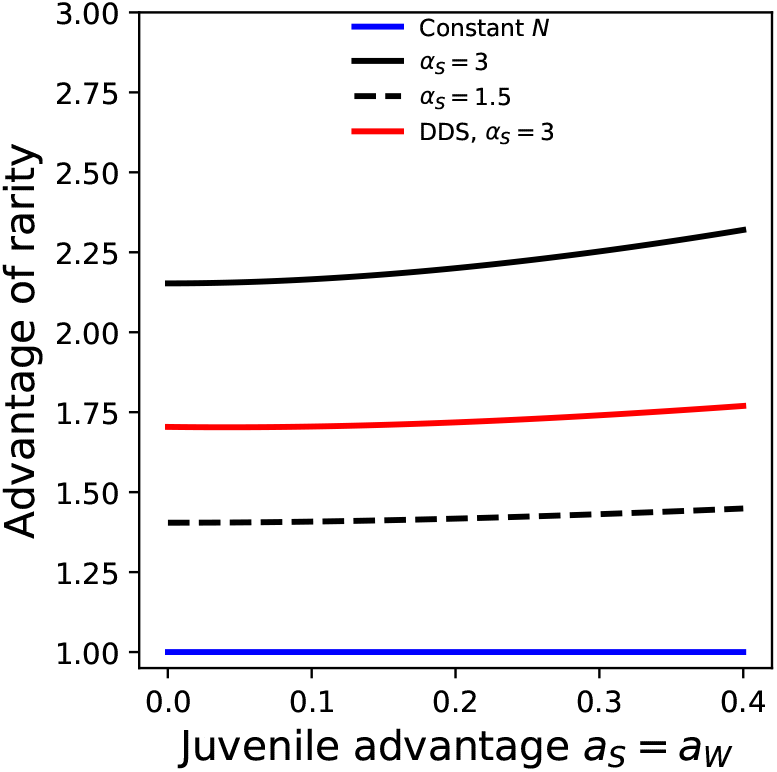
The mere presence of boom-bust cycles enhances the stabilizing effect of protection (solid and dashed black lines; Eq. (12)) compared to the case where N remains constant (blue line), even in the absence of the boom-bust stabilizing mechanism. The red line shows the rarity advantage in Eq. (14), which captures the combined effects of protection, cumulative overdominance, and the boom-bust stabilizing mechanism (this is our more realistic model, in which crowding gradually becomes more important with increasing density and selection is density-dependent). The quantity on the vertical axis is defined in Eq. (13). Simulation parameters: *f* = 0.5, *α_W_* = 0.5, *d_S_* = *d_W_* = 0.5, *i_S_* = *i_W_* = 10, *N*_initial_ = 10^3^, *K* = 10^5^.

Having considered the effects of boom-bust cycles on protection, we now turn to the interaction between the boom-bust stabilizing mechanism and protection. Intuitively, we expect that the combination of protection from selection and boom-bust demography will be “zero-sum” because these mechanisms are most effective under contrasting conditions. Protection is most effective in crowded conditions; protection only inhibits the growth of a common allele if it must displace the rare allele to grow (see “Protection from selection”). On the other hand, the boom-bust stabilizing mechanism is most effective when summer growth starts in uncrowded conditions, since this allows for the largest proportional weakening in the strength of selection due to density-dependence (see “Boom-bust demography”).

Eq. (12) thus exaggerates the stabilizing potential of protection, because juvenile recruitment is modeled as though conditions are crowded at all population densities (the juvenile recruitment rate is divided by the mean relative fitness). To represent the fact that selection at low densities will typically occur in uncrowded conditions, we consider a slightly modified version of Eq. (12) in the summer

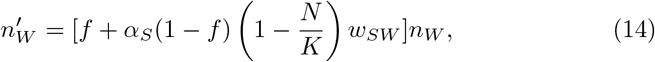

and similarly for a rare summer allele. Here *N* = *n_S_* + *n_W_*, and *K* represents the environmental carrying capacity. In Eq. (14), selection favoring the summer allele gets weaker as density increases (provided that *f* ≠ 0), as required for the boom-bust stabilizing mechanism to work. At low population densities there is no competition and selection is based purely on intrinsic growth rates (*r*-selection). On the other hand, once the population has grown to high density and reached equilibrium, N must be constant which implies that the population mean absolute fitness 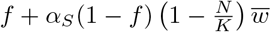 is equal to 1. Substituting this equilibrium requirement into Eq. (14) brings us back to our basic model Eq. (2) again, where selection happens in crowded conditions.

In Fig. 4 we compare the rarity advantages obtained from simulations of Eq. (12) and Eq. (14). It can be seen that, in both of these models, boom-bust demography has a stabilizing effect. The stabilizing effects of protection in Eq. (12) (solid black line) are indeed exaggerated compared to the more realistic treatment of density dependence in Eq. (14) (red line). Nevertheless, the rarity advantage in the latter resulting from the combination of protection in a variable population, cumulative overdominance, and the boom-bust stabilizing mechanism, is still substantially stronger than protection and cumulative overdominance in a constant population (blue line; i.e. the situation summarized by Eq. (4)).

### 3.2 Stabilizing weak versus large effect alleles

In sections 3.1.1–3.1.3 we showed that, for polymorphism to be stable, the difference between *a_S_* and *a_W_* (which represents the extent to which one allele is superior to the other over a seasonal cycle), can be at most first order in the juvenile advantages under reversal of dominance, but at most second order under the other three mechanisms. This has consequences for the stabilization of alleles with weak versus strong fitness effects.

The “effectiveness” of stabilization can be quantified in terms of the greatest difference in juvenile advantages *a_S_* and *a_W_* compatible with stable polymorphism. The simple difference *a_S_* – *a_W_* is not a good measure of effectiveness because the magnitude of this difference will often be proportional to the magnitude of the underlying advantages (the contribution of a locus to fitness sets the scale for *a_S_* and *a_W_*, as well as their difference). Thus, the fact that the regions of stability in Fig. 1 grow wider for larger *a_S_* and *a_W_* does not mean that stabilization is more effective, because the allelic differences that would typically need to be stabilized are growing as well. Consequently, a better measure of effectiveness is the greatest proportional difference compatible with stable polymorphism, given by 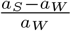 for a rare winter allele, and 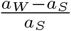 for a rare summer allele.

In the case of reversal of dominance, a rare winter allele can stably segregate with a summer allele with advantage up to 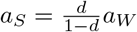, and so the effectiveness of balancing selection is a constant 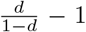. Thus, reversal of dominance is equally effective at stabilizing alleles with weak or strong fitness effects. This results in a broad region of stability even for small juvenile advantages (Fig. 1d).

On the other hand, the effectiveness of balancing selection for cumulative overdominance in Eq. (5) is *d_S_a_S_* (or *d_W_a_W_* for a rare summer allele). Thus, balancing selection is completely ineffective for alleles with very small juvenile advantages, but becomes more effective as the juvenile advantages increase. Similar results apply for protection from selection and boom-bust demography. The combined stability region for cumulative overdominance, protection from selection and boom-bust demography is thus negligibly small for small *a_S_* and *a_W_* (Fig. 1d, red), but rapidly becomes comparable to the reversal of dominance stability region as the juvenile advantages increase. The combination of cumulative overdominance, protection from selection, and boom-bust demogra phy therefore filters out alleles with weak effect while stabilizing stronger effect alleles.

### 3.3 Juvenile advantages and the strength of selection

Cumulative overdominance, protection from selection, and boom-bust demography require *a_S_,a_W_* ~ 0.1 or more to have a meaningful “effectiveness” (see preceding section). Eq. (4) implies an effectiveness of ≈ 10% when the juvenile advantages are ≈ 0.1. The permissible difference *a_W_* – *a_S_* grows rapidly with increasing a*s* thereafter. Such values may seem implausibly high to be prevalent in nature. Even in the temperate *Drosophila* case, where selection at the individual level is regarded as strong, the total change in allele frequencies over a season is only of order ~ 0.1. The number of generations per season is not known (Machado et al., 2018), but each season probably constitutes multiple generations (e.g. Wittmann et al. (2017) suggest ~ 10 generations/season). At face value this does not seem compatible with selection strong enough to allow cumulative overdominance, protection from selection, and boom-bust demography to be effective.

In Fig. 5, we compute the juvenile advantage implied by a given seasonal allele frequency change for an allele at intermediate frequencies. For simplicity, we assume that only protection from selection and cumulative overdominance are present. Since we are considering alleles at intermediate frequencies we use Eq. (1)). Following the temperate *Drosophila* case, we set our given allele frequency change as an increase of 0.1 over the summer from *p_S_* = 0.45 to *p_S_* = 0.55 (most of the fluctuating loci observed by Machado et al. (2018) exhibited smaller fluctuations of 0.04 – 0.08 per season but it is not clear which of these are balanced versus hitchhiking; we follow (Wittmann et al., 2017) in using 0.1 as a representative figure). We consider a range of possible values of summer iterations *i_S_* = 10 – 20. This range of values is based on the assumption that each “season” constitutes half a year, and that the portion of the life cycle from egg laying to reproductive maturity takes ~ 10 – 15 days in *Drosophila* (corresponding to one iteration of our model). Fig. 5 shows that the summer advantage must be at least *a_S_* ~ 0.05 for a seasonal allele frequency change of 0.1. Larger values are needed if there is protection, or if fewer iterations occur per season.

**Figure 5:**
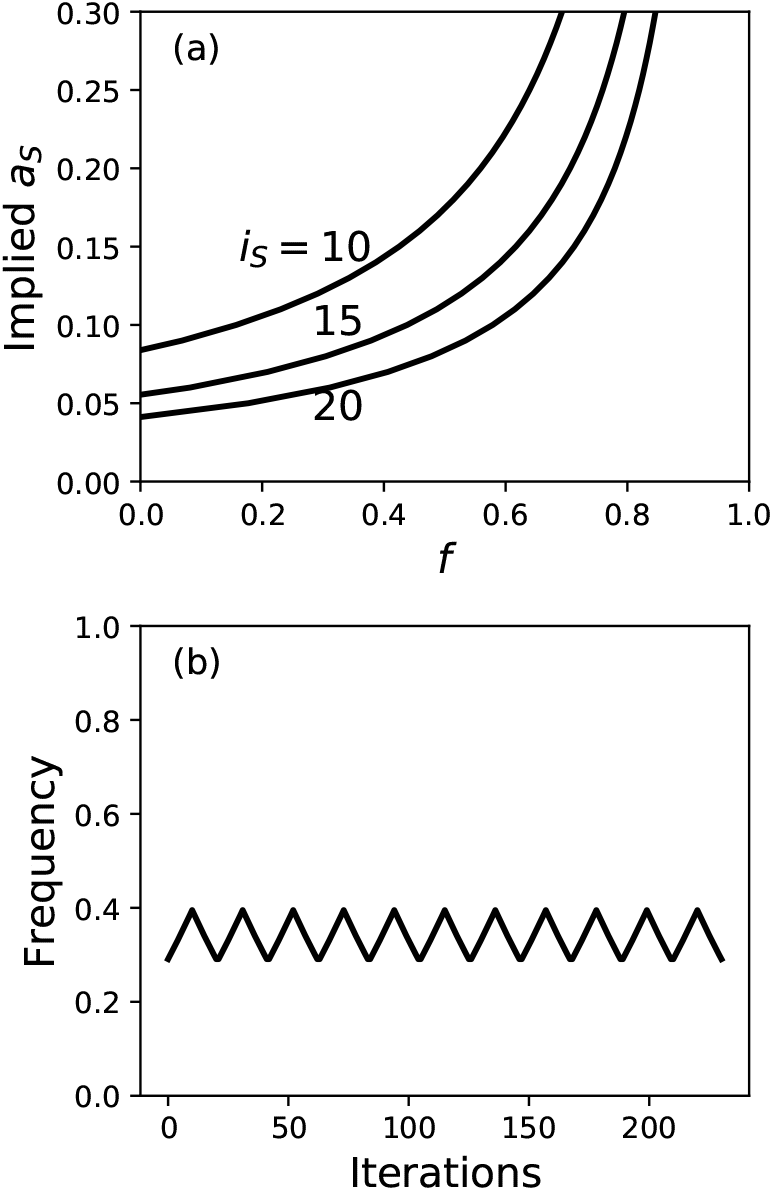
(a) Juvenile advantages of at least ~ 0.05 per iteration are needed to produce a total juvenile allele frequency change of 0.1. Larger values are needed in the presence of protection, or if there are fewer rounds of juvenile recruitment each season. The change in allele frequency is given by Eq. (1). (b) An example of stable polymorphism assuming protection from selection and cumulative overdominance only (*i_S_* = 10, *a_S_* = 0.2, *a_W_* = 0.21, *f* = 0.5, *d_S_* = *d_W_* = 0.5 in Eq. (4)). After running for 100 seasonal cycles to ensure that the steady-state cycle has been reached, the summer allele oscillates between *p* = 0. 3 and *p* = 0. 4.

## 4 Discussion

We have analyzed four different mechanisms that induce balancing selection under temporal variation, the resulting effectiveness of balancing selection, and how these mechanisms interact. Previous analyses critiquing the stabilizing potential of temporal variability considered cumulative overdominance and reversal of dominance only (Smith and Hoekstra, 1980; Hoekstra et al., 1985; Hedrick, 1986), and concluded that only reversal of dominance has a strong enough stabilizing effect to be taken seriously as a basis for balanced polymorphism. Here we also consider the ecological effects of protection from selection and boom-bust demography. In our model, the juvenile advantages a*s* and a*w* determine whether selection is strong enough for balanced polymorphism (Fig. 1). Protection from selection and boom-bust demography expand the region of juvenile advantages compatible with stable polymorphism. Moreover, we show that seasonal allele frequency changes of ~ 0.1, as observed in temperate *Drosophila* populations, imply juvenile advantages of at least ~ 0.05, with larger values needed in the presence of protection (Fig. 5). Together these findings suggest that the scope for temporal variability to balance polymorphism is wider than previous analyses appreciated (Smith and Hoekstra, 1980; Hoekstra et al., 1985; Hedrick, 1986), particularly due to mechanisms other than reversal of dominance.

### 4.4 Empirical tests and implications

#### Protection from selection and cumulative overdominance

The protection from selection mechanism is probably widespread, given some amount of protection from the overlapping generations of iteroparous organisms. However, protection from selection will not be effective at stabilizing polymorphism unless selection is also strong, as measured by the juvenile advantages *a_S_* and *a_W_*. Similar considerations apply for cumulative overdominance; the required incomplete dominance is also likely to be widespread, but will not be effective at stabilizing polymorphism unless selection is strong. The strong selection requirement presumably limits the prevalence of these mechanisms as a stabilizers of polymorphism. However, as we have seen in Sec. 3.3, *Drosophila* data are compatible with juvenile advantages being surprisingly large, particularly if the fraction of protected alleles *f* is appreciable. Specifically, Fig. 5 suggests that juvenile advantages of order ~ 0. 1 are compatible with the magnitude of allele frequency oscillations; this is strong enough for mechanisms other than reversal of dominance to stabilize polymorphism.

Estimating the magnitude of f is challenging. In many models with protection, generational overlap is the primary form of protection, fecundity remains constant with age, and total population density is constant (Chesson and Warner, 1981; Ellner and Hairston, 1994; Svardal et al., 2015). The strength of protection f is then simply the fraction of surviving adults each round of juvenile recruitment. With this simple model model in mind we can get a rough estimate of the magnitude of *f* in wild *Drosophila* from laboratory measurements. Female fecundity peaks ≈ 5 days after reaching reproductive maturity but egg laying continues up to 20 days after which female survivorship declines rapidly (Le Bourg and Moreau, 2014). We approximate this as death occurring at 20 days. Assuming that it takes 10 days to develop from an egg to reproductive maturity, this means that females persist for roughly two rounds of juvenile recruitment under laboratory conditions. This corresponds to a protection due to overlapping generations of *f* = 0.5. This number is probably an overestimate since females in the wild are subject to additional sources of mortality such as predation and parasitism, and female fecundity declines with age.

In reality, protection is more complicated. Adults may also experience some selection, e.g. due to predation/parasites. In this case, protected alleles only experience weakened negative selection, not complete protection; 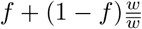 in Eq. (2) would then be replaced by 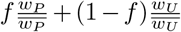 where *P* and *U* stand for “protected” and “unprotected”, respectively. Additional complications arise in an expanding population. For instance, the strength of protection may depend on how rapidly the population is growing; even if the proportion of surviving adults is large, they will only constitute a small fraction of the population if a large number of juveniles is recruited to adulthood each iteration. Other forms of protection include maternal effects, which can confer protection because, in any given iteration, some fraction of unfavored alleles will have a favored mother (e.g. the heterozygote offspring of a summer homozygote in the summer). We would then need to distinguish between the possible parental combinations that an allele might experience (Wolf and Wade, 2016). The development of population genetic methods to quantify protection from selection is needed.

Finally, note that protection depends upon the presence of crowding (section “Combining protection and boom-bust demography”). While crowding is likely to be important in organisms with relatively stable demography, which are presumably at or near their environmental carrying capacity, the presence of strong crowding effects is less certain in populations which undergo cyclical collapses in population density.

#### Boom-bust demography

Boom-bust demography is distinguished by its reliance on density-dependent selection induced by crowding. A relatively simple way to test for the presence of stabilization driven by boom-bust demography would be to alleviate the effects of crowding by artificially constraining density e.g. via culling of adults. Adult culling would also mitigate the effects of overlapping generations (which would also be affected by changes to crowding). If boom-bust demography is indeed stabilizing polymorphism, we would then expect temporally-averaged allele frequencies to shift (potentially even losing the polymorphism), as a result of the weakening of the boom-bust demography rarity advantage.

#### Reversal of dominance

Reversal of dominance is distinguished by its high effectiveness: it is an order of magnitude stronger than cumulative overdominance, protection from selection and boom-bust demography. Among the mechanisms considered here, reversal of dominance is unique in being insensitive to the magnitude of the juvenile advantages (Sec. 3.2). Thus, we might be able to test for the presence of reversal of dominance stabilization by experimentally weakening the juvenile advantages *a_S_* and *a_W_* by similar proportions. We would then expect to only see a reduction in the amplitude of allele frequency oscillations, with comparatively little change in temporally-averaged allele frequencies. The reason for this is that the temporally-averaged allele frequency is set by a balance between the superior juvenile advantage of one allele and the rarity advantage of the other; under reversal of dominance these both scale with the juvenile advantage. By contrast, under cumulative overdominance, protection from selection and boom-bust demography, the rarity advantage rapidly disappears for smaller *a_S_* and *a_W_*, and we would expect to see the allelic superiority overwhelming it if we weaken selection.

#### Multi-locus polymorphism

Our analysis has considered the simple case of two alleles segregating at a single locus. To understand pervasive balancing selection we must confront the multi-locus case (Wittmann et al., 2017), which requires us to account for the interactions between loci. These interactions can occur via linkage or epistasis. Note that, even if recombination rates are high, overlapping generations in the presence of selection will generate linkage disequilibrium. Linkage can either act to preserve or eliminate genetic variation at a given locus depending on whether nearby loci are under directional or balancing selection. As such, linkage does not directly promote or eliminate balanced polymorphism (although linkage can stabilize polymorphism in combination with sign epistasis; Novak and Barton 2017).

Many different forms of epistasis are possible, some of which can dramatically alter the stability of polymorphism at individual loci (Novak and Barton, 2017; Gulisija et al., 2016). However, at the genomic scale there is empirical support for diminishing returns epistasis when combining the beneficial effects of segregating alleles across loci (Chou et al., 2011; Kryazhimskiy et al., 2014). Diminishing returns epistasis has important consequences for multi-locus polymorphism if weak effect alleles can be stabilized: weak effect polymorphic loci will accumulate and preclude larger effect polymorphisms from being present in mutation-selection-drift balance as a result of diminishing returns (Wittmann et al., 2017). This poses a problem for reversal of dominance, but not cumulative overdominance, protection from selection and boom-bust demography (Sec. 3.2).

We can therefore make the following conclusions about multi-locus polymorphism in *Drosophila.* If reversal of dominance is indeed responsible for balancing these polymorphisms, then it must somehow get around its weak allele problem. One way this might occur is if reversing dominance is less likely to occur in weak effect alleles. Unless there is a solution to the weak allele problem, mechanisms which do not stabilize weak effect alleles must be responsible for stabilizing many of the observed polymorphisms (assuming that these alleles are indeed balanced; migration is one important alternative); this could include the three weaker mechanisms considered here. Although we have shown that seasonally oscillating selection in *Drosophila* could plausibly be strong enough for these weaker mechanisms to be effective, a more definitive conclusion rests upon either estimating the magnitude of f, or on direct measurements of the juvenile advantages *a_S_* and *a_W_*.

### 4.2 Extensions and limitations

Much of our analysis rests on standard population genetic models of selection in diploids, and is therefore subject to the usual caveats (e.g. the lack of frequency-dependence in *a_S_* and *a_W_*, nonrandom mating, and so on). Our extension of these standard models to incorporate protection is derived from classical models of protection induced by generational overlap (Chesson and Warner, 1981; Svardal et al., 2015). As discussed above, our description of protection would need to be modified to describe more complicated forms of protection.

In our treatment of boom-bust demography we have only considered one model that explicitly represents density-dependent selection (Eq. (14)). Many other choices are possible here, which could depend on the model of protection. The variable-density territorial contest model of Bertram and Masel (2019), which has overlapping generations as its protection mechanism, is one possibility for exploring some of these issues. One important possibility that we have not discussed is if density dependence causes selection to switch sign such that the winter type is favored at high density (e.g. due to a growth/competitive ability trade-off). We have specifically avoided this scenario here since it would allow polymorphism in a stable environment and our focus is on fluctuation-dependent stability mechanisms.

Due to our focus on fluctuation-dependent stability mechanisms, we have also not attempted to explore spatial variation or phenotypic variation within genotypes (individual variation), or how different sources of variation interact (see Frank and Slatkin (1990) and Svardal et al. (2015)).

When evaluating whether two alleles will stably persist, we have taken the allelic juvenile advantages to be fixed quantities. In the long term the juvenile advantages will evolve, leaving open the possibility that polymorphism is not evolutionarily stable (Ellner and Hairston, 1994). This scenario is addressed by Svardal et al. (2015), who derive conditions for polymorphism in variable environments based on “evolutionary branching”. That is, Svardal et al. (2015) require a rarity advantage at an attractor in the dynamics of a quantitative trait evolving by sequential substitutions of small effect mutations. Our model does not have any attractors in this sense, because in the absence of further assumptions, evolution will favor generalist alleles with ever increasing advantages in both seasons. This is not biologically reasonable because it fails to account for trade-offs in traits associated with summer versus winter success. In the absence of a model of these trade-offs, it is therefore not possible to further connect the findings of Svardal et al. (2015) to ours.

This points to a broader divide between population genetic and more phenotypic approaches (such as adaptive dynamics) to the study of genetic variation. Adaptive dynamics approaches typically view variation through the lens of evolutionary branching, which implies disruptive selection at the phenotypic level. One consequence of disruptive selection is that fitness-associated genetic variation is expected to become concentrated at a few loci, not dispersed over many, because recombination is more likely to produce less fit intermediate phenotypes when genetic variation is dispersed (Kopp and Hermisson, 2006; van Doorn and Dieckmann, 2006; Svardal et al., 2015). The population genetic approach does not require phenotypic disruptive selection, and predicts no tendency for genetic variation to be concentrated (Wittmann et al., 2017). However, population genetic approaches generally do not specify the phenotypic basis of selection at all. This quickly becomes problematic when trying to account for ecological factors, as we have seen in the case of boom-bust demography. There is therefore a need to better integrate the population genetic and phenotypic approaches.

## Appendix A: Details of the diploidy and protection from selection conditions

Here derive the condition for stable polymorphism under the combined effects of diploid Mendelian inheritance and protection from selection, Eq. (8). Recall that this condition assumes *i_S_* = *i_W_*, and so we can take the *i_W_* ‘th root to remove the exponents in Eq. (3). Multiplying by *w_SS_*(*S*)*w_SS_*(*W*), Eq. (3) can thus be written as

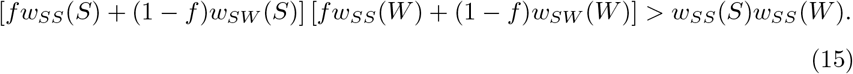

Substituting from Table 1 we have,

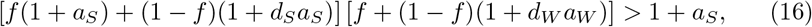

which can be simplified to

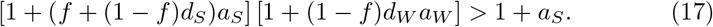

Multiplying out the square brackets and subtracting the right hand side gives

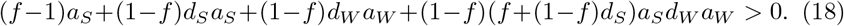

Dividing by (1 – *f*) then gives

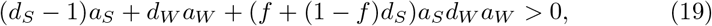

which yields Eq. (4).

## Appendix B: Details of the boom-bust condition

Here we derive Eq. (10), which gives gives the condition for stable polymorphism in the presence of boom-bust population demography assuming a simplified representation of density-dependence in which only r-selection occurs (see section “Boom-bust demography” for assumptions and notation).

If the winter allele is rare, the summer homozygote determines the overall rate of population expansion, and we have (*b_SS_*)^*i*_*S*|Wrare_^ = *D*, or equivalently *i*_*s*|Wrare_ = ln *D*/ln *b_SS_*. On the other hand, if the summer allele is rare, then *i*_*S*|Srare_ = ln *D*/ln*b_WW_* = ln *D*/[ln *b_SS_* – ln(*b_SS_*/*b_WW_*)], where *b_SS_*/*b_WW_* = *w_SS_*(*S*)/*w_WW_* (*S*) = 1 + *a_S_*. Thus, to first order in *a_S_* we have ln *b_SS_*/*b_WW_* ≈ *a_S_*, and

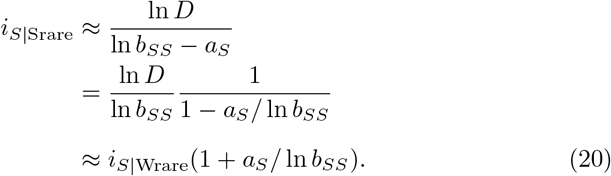

Defining *i* = (*i*_*S*|Srare_ + *i*_*S*|Wrare_)/2 as the “baseline” number of summer iterations, we therefore have *i*_*S*|Srare_ ≈ *i*(1 + *ca_S_*) and *i*_*s*|Wrare_ ≈ *i*(1 – *ca_S_*) to first order in *a_S_*, where *c* = 1/(2ln *b_SS_*).

The strength of the boom-bust stabilizing mechanism is controlled by the constant *c* = 1/(2ln *b_SS_*). This dependence of *c* on *b_SS_* is a consequence of the discrete-time nature of our model. In the continuous time limit where each iteration represents a vanishingly small growth increment, *c* → 1/2 and the boom-bust stability mechanism depends entirely on relative ratios between absolute Malthusian parameters (compare Yi and Dean 2013). In practice, c will typically be of close to 1/2 unless growth is extremely rapid (e.g. *c* ≈ 0.72 in a population which doubles every summer iteration). We therefore set *c* =1/2 in Fig. 1.

For simplicity, in Eq. (9) we assume symmetry between the seasons, such that the winter has the same number of iterations as the baseline number of summer iterations, i.e. *i_W_* = *i* with is = *i_W_* in the case where *a_S_* = 0. In the asymmetric case where either summer or winter is intrinsically longer, a different region of juvenile advantages can permit stable polymorphism. For instance, the summer advantage must be larger for a given winter advantage if *i_W_* > *i*. But this asymmetry does not affect the overall magnitude of the stabilized region, because constant differences between *i_S_* and *i_W_* do not introduce frequency dependence.

Taking the *i_W_*‘th root of Eq. (3) with *f* = 0, our stability condition for a rare winter allele can thus be written analogously to Eq. (17) as

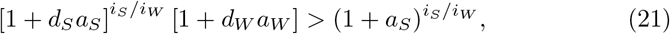

where *i_S_/i_W_* = 1 – *ca_S_*. We then apply the generalized binomial theorem: 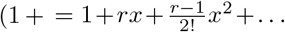 This implies 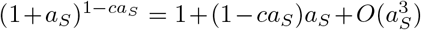, and similarly 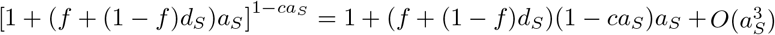. Neglecting third order terms in *a_S_*, Eq. (21) is thus the same as Eq. (17), but with *a_S_* replaced by (1 – *ca_S_*)*a_S_* and *f* = 0. The same steps used to obtain Eq. (4) from Eq. (17) (Appendix A) will therefore yield Eq. (4) with *a_S_* replaced by (1 – *ca_S_*)*a_S_*. In the resulting inequality, the *ca_S_* contribution on the right hand side is 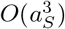 and can be neglected, and we move the *ca_S_* contribution on the left hand side to the right to obtain Eq. (10).

The corresponding condition for a rare summer allele is obtained by swapping *S* and *W* labels in Eq. (17) and then replacing *a_S_* → (1– *ca_S_*)*a_S_*, again neglecting 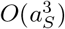 terms and setting *f* = 0. The *ca_S_* contribution on the left now appears in the term –*d_S_*(1 – *ca_S_*)*a_S_*; the boom-bust demography term on the right is thus 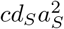 for a rare summer allele.

## Appendix C: Protection in the presence of boom-bust cycles

Here we give some mathematical details of how protection behaves in the presence of boom-bust cycles. Applying the same procedure as outlined in Appendix A to the haploid version of Eq. (12), it can be shown that the condition for stable polymorphism is given by

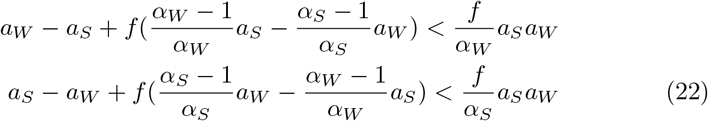

for the rare-*S* and rare-*W* cases respectively. The last term on the left represents the fact that the juvenile advantages *a_S_* and *a_W_* no longer contribute on equal terms to per-capita growth due to the asymmetrical effects of protection in the summer and winter. This term simply switches sign when the *S* and *W* labels are switched in the rare-*W* versus rare-*S* cases, and it therefore makes zero contribution to the overall advantage of rarity in Eq. (13). The enhancement of the overall region of stability arises due to the fact that in a population undergoing boom-bust cycles, 1/*α_W_* ≫ 1 on the right hand side. Provided that the boom-bust cycles are large enough (i.e. *α_W_* is small enough), this will more than offset the loss of coexistence due to the fact that 1/*α_S_* < 1 in the rare-*W* case – but the region of coexistence will now be asymmetric.

Contributions
JB and JM conceived the project. JB performed the analysis. JB and JM wrote the manuscript.

## Acknowledgements

We thank Meike Wittmann, Dmitri Petrov and Michael Wade for helpful discussions, and Peter Chesson for comments on an earlier version of the manuscript.

